# Mast-Cell Expressed Membrane Protein-1 (MCEMP1) is expressed in classical monocytes and alveolar macrophages in Idiopathic Pulmonary Fibrosis and regulates cell chemotaxis, adhesion, and migration in a TGFβ dependent manner

**DOI:** 10.1101/2023.10.07.561349

**Authors:** Carole Y. Perrot, Theodoros Karampitsakos, Avraham Unterman, Taylor Adams, Krystin Marlin, Alyssa Arsenault, Amy Zhao, Naftali Kaminski, Gundars Katlaps, Kapilkumar Patel, Debabrata Bandyopadhyay, Jose D. Herazo-Maya

**Affiliations:** Division of Pulmonary, Critical Care and Sleep Medicine, Ubben Center for Pulmonary Fibrosis Research, Department of Internal Medicine, Morsani College of Medicine, University of South Florida, Tampa, FL, USA; Section of Pulmonary, Critical Care and Sleep Medicine, Department of Internal Medicine, School of Medicine, Yale University, New Haven, CT, USA; Pulmonary Fibrosis Center of Excellence, Institute of Pulmonary Medicine, Tel Aviv Sourasky Medical Center, Sackler School of Medicine, Tel Aviv University, Tel Aviv, Israel; Genomic Research Laboratory for Lung Fibrosis, Tel Aviv Sourasky Medical Center, Tel Aviv, Israel; Division of Cardiothoracic Surgery, Department of Surgery, Morsani College of Medicine, University of South Florida, Tampa, FL, USA; Center for Advanced Lung Disease and Lung Transplant Program. Tampa General Hospital, FL, USA

**Author notes:** Correspondence to: Jose D. Herazo-Maya, MD, Ubben Endowed Chair and Associate Professor of Medicine Chief of Pulmonary, Critical Care and Sleep Medicine, Director of the Ubben Center for Pulmonary Fibrosis Research Department of Internal Medicine, Morsani College of Medicine University of South Florida. Equal contribution (Carole Y. Perrot, Theodoros Karampitsakos).

**Keywords:** Mast-Cell Expressed Membrane Protein-1, monocytes, alveolar macrophages, Idiopathic Pulmonary Fibrosis

## Abstract

**Background:** Mast-Cell Expressed Membrane Protein-1 (MCEMP1) is higher in Idiopathic Pulmonary Fibrosis (IPF) patients with increased risk of death and poor outcomes. Here we seek to establish the mechanistic role of MCEMP1 in pulmonary fibrosis.

**Methods:** MCEMP1 expression was analyzed by single-cell RNA sequencing, immunofluorescence in Peripheral Blood Mononuclear Cells (PBMC) as well as in lung tissues from IPF patients and controls. Chromatin Immunoprecipitation (ChiP) and Proximity Ligation Assay (PLA) were used to study the transcriptional regulation of *MCEMP1*. Transient RNA interference and lentivirus transduction were used to knockdown and knock-in MCEMP1 in THP-1 cells to study chemotaxis, adhesion, and migration. Bulk RNA sequencing was used to identify the mechanisms by which MCEMP1 participates in monocyte function. Active RHO pull-down assay was used to validate bulk RNA sequencing results.

**Results:** We identified increased MCEMP1 expression in classical monocytes and alveolar macrophages in IPF compared to controls. MCEMP1 was upregulated by TGFβ at the mRNA and protein levels in THP-1. TGFβ-mediated MCEMP1 upregulation results from the cooperation of SMAD3 and SP1 via concomitant binding to SMAD3/SP1 *cis*-regulatory elements within the *MCEMP1* promoter. In terms of its function, we found that MCEMP1 regulates TGFβ-mediated monocyte chemotaxis, adhesion, and migration. 400 differentially expressed genes were found to increase after TGFβ stimulation of THP-1, further increased in MCEMP1 knock-in cells treated with TGFβ and decreased in MCEMP1 knockdown cells treated with TGFβ. GO annotation analysis of these genes showed enrichment for positive regulation of RHO GTPase activity and signal transduction. While TGFβ enhanced RHO GTPase activity in THP-1 cells, this effect was attenuated following MCEMP1 knockdown.

**Conclusion:** MCEMP1 is highly expressed in circulating classical monocytes and alveolar macrophages in IPF. MCEMP1 is regulated by TGFβ and participates in the chemotaxis, adhesion, and migration of circulating monocytes by modulating the effect of TGFβ in RHO activity. Our results suggest that MCEMP1 may regulate the migration and transition of monocytes to monocyte-derived alveolar macrophages during pulmonary fibrosis development and progression.

## Introduction

Idiopathic pulmonary fibrosis (IPF) is a chronic and progressive interstitial lung disease (ILD) characterized by repetitive epithelial cell injury, aberrant deposition of extracellular matrix, immune deregulation, irreversible lung tissue scarring and subsequently impaired lung function (1). Current antifibrotic therapies are only able to slow lung function decline (2, 3). Thus, disease progression is inevitable. However, the pattern of disease progression is highly heterogeneous, with some patients demonstrating long term clinical stability and others experiencing a more rapid disease course. This fueled extensive research effort for the identification of biomarkers predictive of IPF progression and mortality (4).

We previously used genome wide, transcript profiling of peripheral blood mononuclear cells to identify expression patterns predictive of IPF survival and validated a peripheral blood 52-gene signature predictive of transplant-free survival and mortality in IPF patients from six different cohorts (5, 6). We have previously shown by cellular deconvolution and single-cell RNA sequencing that classical monocytes in peripheral blood are the source of increased gene expressions predictive of mortality in IPF (7). Mast-cell expressed membrane protein 1 (MCEMP1, c19orf59) a novel and poorly characterized gene, was among the upregulated genes of the 52-gene signature (6). The *MCEMP1* gene encodes a type II transmembrane protein and its expression has been identified in monocytic leukemia cell lines (THP-1) and in lung mast cells (8).

While we studied peripheral blood gene expression changes predictive of IPF mortality, we did not study the cellular source and the mechanisms of MCEMP1 in the pathogenesis of pulmonary fibrosis. In this study, we showed that MCEMP1 is expressed in circulating monocytes and in alveolar macrophages in IPF. We also identified that MCEMP1 is transcriptionally regulated by TGFβ in a SMAD3/SP1-dependent manner. Our study shows that MCEMP1 might modulate TGFβ-mediated monocyte adhesion, chemotaxis and migration by regulating RHO-GTPase activity. Our results demonstrate that *MCEMP1* is a TGFβ target gene that may be critical in the migration and transition of circulating monocytes to monocyte-derived alveolar macrophages during lung injury and aberrant repair.

## Material and Methods

### Single-cell RNA sequencing data analysis

We re-analyzed peripheral blood mononuclear cell single-cell RNA sequencing (scRNA-seq) data from N=25 IPF patients and N=13 controls recruited at Yale University as previously reported (9). Comparison of MCEMP1 expression levels between monocytes of IPF patients and controls was conducted using the Wilcoxon rank sum test. We also re-analyzed single-cell RNA sequencing dataset including 312,928 cells from distal lung parenchyma samples obtained from 32 IPF lungs (45 libraries yielding 147,169 cells), 18 chronic obstructive pulmonary disease (COPD) lungs (24 libraries yielding 69,456 cells), and 28 control donor lungs (38 libraries, yielding 96,303 cells)(10).

### Detection of CD206, MCEMP1 and HT2-280 in human lungs by immunofluorescence staining and quantification

Normal lung tissue slides were purchased from Origene (Rockville, MD) and used as controls. IPF lung tissues were explants derived from three patients undergoing lung transplantation at Tampa General Hospital. Briefly, fresh human lung tissue was collected, placed on ice-chilled Petri dishes, and cut into 5 mm thick-pieces. Lung tissues were washed twice in 1X PBS, fixed in 10% formalin overnight at 4°C on a rocker, progressively dehydrated in 70%, 80%, 90%, 95% and 100% ethanol, cleared in xylene (2x 30 min) and embedded in paraffin (3x 60 min) using a Leica Automatic Tissue Processor (Wetzlar, Germany). Paraffin tissue blocks were left overnight at room temperature to solidify prior sectioning. Sample collection was approved by the local IRB board. For immunofluorescence staining, tissue sections were deparaffinized, hydrated, immersed in a pH=6 antigen retrieval solution (IHC World, Elliott City, MD) and placed in a steamer for 40 min at 95-98°C. Slides were allowed to cool down for 20 min, then rinsed with water and air-dried completely.

Tissue sections were incubated in blocking buffer (Triton X-100 0.1%, goat serum 10%, BSA 3%, 1X PBS) for an hour at room temperature, followed by an overnight incubation with different combinations of primary antibodies targeting CD206 (5ug/ml), MCEMP1 (1.5ug/ml), and type-2 alveolar epithelial cell marker HT2-280 (1/200) at 4°C. Tissue sections were washed three times with 1X PBS and incubated with combinations of Alexa-Fluor 488, 555 and 647 secondary antibodies (1/1000 dilution; ThermoFisher scientific) for one hour at room temperature followed by DAPI for 5 min. Tissue sections were washed three times with 1X PBS, mounted under coverslip using Cytoseal 60, and stored at 4°C overnight before image capture using a Nikon Eclipse Ni-E fluorescence microscope and NIS-Elements software.

MCEMP1 and CD206 staining were quantified using NIS Elements AR5-30-05 software. Macrophages, identified as CD206^+^ cells, and MCEMP1^+^ cells were quantified by measuring the ratio CD206 ^+^ area/total tissue area and MCEMP1 ^+^ area/total tissue area, respectively, in >20 micrographs from two healthy and three IPF lungs. MCEMP1 ^+^ macrophages were quantified by measuring MCEMP1^+^; CD206 ^+^ cells/total CD206^+^ cells. The proportion of macrophages among MCEMP1^+^ cells was measured with the ratio MCEMP1 ^+^; CD206 ^+^ cells/total MCEMP1 ^+^ cells. Statistical analysis was performed using One-way ANOVA with Sidak post-hoc test for multiple comparisons.

### Cell culture and reagents

THP-1 and U937 cell lines (Millipore Sigma, Burlington, MA) were cultured in RPMI 1640 medium with L-glutamine (Gibco, ThermoFisher Scientific, Waltham, MA) supplemented with 10% fetal bovine serum and antibiotics. All our experiments were performed using THP-1 and U937 at passage <25. Recombinant human TGFβ1, FGF2, TNF alpha, IL14, IL10 and IL14 (Peprotech, Cranbury, NJ) and LPS (ThermoFisher) were used to stimulate THP-1 cells. Phorbol 12-myristate 13-acetate (PMA) was used for monocyte to macrophage differentiation. SMAD3 inhibitor SiS3 and HDAC inhibitor trichostatin A were purchased from Millipore Sigma and Mithramycin A from Cayman Chemical (Ann Harbor, MI).

### Antibodies

For western blot and immunostaining applications, we used a polyclonal rabbit anti-MCEMP1 antibody from Millipore Sigma (ref. HPA014731). For western blot only, monoclonal rabbit anti-SMAD3 (ref. 9523), anti-phospho-SMAD3 (ref. 9520), anti-SMAD2 (ref. 5339), anti-phospho-SMAD2 (ref. 18338), and anti-GAPDH (ref. 5174) antibodies were purchased from Cell Signaling Technology (Danvers, MA). All secondary HRP-conjugated antibodies were obtained from Promega (Madison, WI). For immunofluorescence only, monoclonal mouse anti-CD206 (MRC1; ref. Amab90746) was from Millipore Sigma, monoclonal mouse anti-CD14 (ref. MA-1-23611) was from Invitrogen (ThermoFisher Scientific) and monoclonal mouse anti-HT2-280 (ref. T-27) was from Terrace Biotech (San Francisco, CA). Alexa Fluor 647 Phalloidin as well as Alexa Fluor secondary antibodies (488, 555, and 647) were purchased from ThermoFisher Scientific. For chromatin immunoprecipitation, we used ChIP grade monoclonal rabbit anti-SP1 (ref. ab231778) and anti-SMAD3 (ref. ab208182) antibodies from Abcam (Cambridge, MA). For proximity ligation assay, we used the anti-SP1 antibody mentioned above and a monoclonal mouse anti-SMAD3 antibody from Proteintech (Rosemont, IL).

### RNA extraction and RT-qPCR

RNA extraction was conducted using Rneasy Plus kit from Qiagen (Hilden, Germany) following the manufacturer’s instructions. Between 500 ng and 1 µg of RNA was then subjected to Dnase I, Amplification Grade digestion (ThermoFisher Scientific) followed by reverse transcription using Superscript IV First-Strand Synthesis System (ThermoFisher Scientific). Quantitative polymerase chain reaction (qPCR) was then performed using Power SYBR Green PCR Master Mix (ThermoFisher Scientific) on a CFX384 Real-Time PCR detection system (Biorad. Hercules, CA). Primer sets for MCEMP1, RPS18 and RPL37A were designed using Primer Blast (11) and empirically validated for quantitative PCR use following primer efficiency calculation. All our qPCR experiments were repeated at least three times, and samples were run in triplicates.

### Protein isolation and western blot

Total cell proteins were extracted using radioimmunoprecipitation (RIPA) buffer supplemented with protease inhibitor cocktail and phosphatase inhibitors, and BCA assay was performed to determine protein concentration (ThermoFisher). Western blotting was performed by electrophoresis of 20 µg of proteins on Mini-PROTEAN or Criterion TGX Precast Gels (Bio-Rad, Hercules, CA, USA), followed by electrotransfer to Amersham™ Protran® nitrocellulose membrane (Millipore Sigma). After blocking unspecific binding, antibody incubations were carried out overnight in blocking buffer (5% BSA or 5% nonfat milk in TBS containing 0.1% Tween-20), and target proteins were detected using Western Lightning Plus-ECL (PerkinElmer, Waltham, MA) and a Chemidoc imaging system (Biorad). Blots were quantified using ImageJ. All our WB experiments were repeated at least three times, and samples were run in triplicates.

### Live Cell Imaging

THP-1 cells were seeded in 6 well-plates and treated with TGFβ (5ng/ml) for 48 hours, followed by an overnight incubation with PMA (150nM). Cells were washed three times with 1X PBS and incubated with MCEMP1 (1/200) or CD14 (1/200) antibodies diluted in ice-cold assay buffer (1X PBS, BSA 2%, NaN _3_ 0.5%) for 2 hours at 4 °C. After two more washes with the assay buffer, cells were incubated with Alexa Fluor 555 secondary antibodies for 1.5 hour at 4 °C, washed twice with the assay buffer and resuspended in PBS for observation and picture acquisition using a Nikon fluorescence microscope.

### Chromatin immunoprecipitation

THP-1 cells were grown in T75 flasks for 24 hours, then treated with SiS3 (25 µM), mithramycin A (100 nM), trichostatin A (10 µM) or vehicle for one hour before TGFβ treatment. Twenty-four hours later, cells were harvested, spun down at 300g for 5 min and cell pellets were resuspended in 1% formaldehyde fixation solution for 12 minutes at room temperature under gentle agitation. Chromatin immunoprecipitation (ChIP) was carried out using the ChIP-IT Express Enzymatic kit from Active Motif (Rixensart, Belgium). In brief, 10 μg of enzymatically sheared chromatin was incubated overnight at 4°C using 3 μg of either control anti-IgG, ChIP grade anti-SMAD3 (Abcam), or ChIP grade anti-SP1 (Abcam) antibodies along with protein G magnetic beads. Precipitated chromatin was eluted, cross-links reversed, submitted to proteinase K treatment, and processed for quantitative PCR analysis of SMAD3 and SP1 binding to the human MCEMP1 promoter using Power SYBR Green PCR Master Mix (ThermoFisher Scientific). ChIP-qPCR results were calculated using the ΔΔCt method and are shown as percentage of input DNA for each ChIP experiment.

### Proximity Ligation Assay

The DuoLink *in Situ* Detection reagents Orange (λex = 554 nm and λem = 576 nm; Millipore Sigma) was used to determine whether SMAD3 interacts with SP1 in THP-1 cells. Briefly, PMA-treated cells were grown in 4-well chamber slides until attachment, then subjected to SiS3, mithramycin A or vehicle treatment followed by TGFβ for 24 hours. Cells were washed with 1x PBS, fixed with a 4% paraformaldehyde solution for 10 min, permeabilized for 40 min at room temperature, blocked with the Duolink® Blocking Solution for one hour at 37°C, and incubated overnight at 4°C with mouse anti-SMAD3 (Proteintech, 1:400) and rabbit anti-SP1 (Abcam, 1:200) antibodies. Samples were then incubated with DNA-conjugated secondary antibodies (anti-rabbit probe PLUS and anti-mouse probe MINUS, 1:5 dilution) at 37°C for 1 h. Our negative control consisted in a sample incubated with secondary antibodies only. Hybridization, ligation, amplification, and detection steps were performed according to the manufacturers’ instructions. Cover slips were mounted on slides with mounting medium with 40,6-diamidino-2-phenylindole (DAPI) from the kit and kept at −20°C overnight before image capture and analysis using a Nikon Eclipse Ni-E fluorescence microscope and NIS-Elements software.

### Transient RNA interference

For MCEMP1 silencing, THP-1 cells were transfected at 600,000 cells/ml in RPMI 10% FBS with 10 nM Silencer Select siRNA (Ambion; sense 5’GAAUGUCUCAAACUCCGUAtt – antisense 5’UACGGAGUUUGAGACAUUCca) using Lipofectamine RNAiMAX Reagent (ThermoFisher Scientific). Cells were collected 48- and 72-hours post-transfection for analyses.

### Lentivirus transduction

For MCEMP1 stable silencing and overexpression, THP-1 cells were transduced by spinofection with either pGFP-C-shMCEMP1 or -shctrl lentiviral vectors for silencing, and either with a lentiviral expression vector carrying GFP-tagged MCEMP1 ORF or insertless (mock) vector for MCEMP1 overexpression (lentivirus were purchased from Origene, MD, USA). A MOI of 50 was necessary to achieve a satisfactory infection rate. Transduced cells were selected with puromycin (0.5μg/ml) for 10 to 15 days and tested for MCEMP1 expression by western blot before use. All primer sets used for our study can be seen in Table S1.

### Chemotaxis assay

The Chemotaxis assay was performed using the Cell Migration/Chemotaxis Assay kit (24-well, 5μm) from Abcam following the manufacturer’s instructions. Briefly, we performed transfection of THP-1 cells with control- or MCEMP1-targeting siRNA to knockdown MCEMP1 expression, followed by TGFβ stimulation for 24 hours. We also stably transduced THP-1 cells with mock- and MCEMP1-lentiviral vectors to induce MCEMP1 expression. Cells from these experimental conditions were collected, washed with 1X PBS, resuspended in basal RPMI and 200,000 cells were seeded into 5um-pore sized inserts of 24 well-transwell plates. Serum-free medium (0.1% FBS) containing chemoattractant CCL2 (50ng/ml) was dispensed in the bottom chambers of the plates, and cells were incubated at 37 °C for 18 to 24 hours. Inserts were carefully aspired and removed, and the plates were centrifuged at 1000g for 5 min. The number of migrated cells was determined by measuring the fluorescence signal intensity for each sample using a standard curve.

### Cell adhesion assay

THP-1 transfected either with control- or MCEMP1-targeting siRNA were cultured in 6-well plates and stimulated with TGFβ (5ng/ml) 24 hours post-transfection. Cells were collected 24 hours later, washed with 1X PBS, resuspended in serum-free medium and seeded in 24 well-plates (200,000 cells/well). After incubation for one hour at 37°C, unattached cells were removed with PBS and adherent cells were fixed with a 20% methanol/0.5% crystal violet solution for 15 min at room temperature. PMA-treated cells were used as positive control. Crystal violet was extracted from cells using 100 μl of ethanol 70% and absorbance was measured at 595 nm. The assay was repeated three times in tri-or quadruplicates for each condition.

### Wound closure (scratch) assay

Mock- and MCEMP1-overexpressing THP-1 cells (from two independent transduction experiments) were seeded in 6-well plates at the density of 1 x 10 ^6^ cells/dish and immediately treated with 150 nM PMA to induce THP-1 differentiation into macrophages and cell adhesion. Forty-eight hours post-treatment, a scratch was performed with a 200 μl-pipette tip to create a wound in the confluent cell monolayer. Wound closure was monitored by taking bright field pictures of the cells immediately after scratch and 72 h later using optical microscopy. The ImageJ software was used to measure wound closure areas. The experiment was performed in two independent wells for each condition and our results represent the analysis of 10 micrographs.

### Bulk RNA sequencing and analysis

To study the mechanisms by which MCEMP1 participates in monocyte chemotaxis, we performed bulk RNA-sequencing using three batches of transduced cells (see lentivirus transduction section above) that were sub cultured and treated with TGFβ (5ng/ml) or vehicle for 24 hours before RNA extraction. RNA-seq was performed by Genewiz-Azenta Life Sciences (South Plainfield, NJ, USA). Differential gene expression analysis was performed using Deseq2 package. Three groups were analyzed: non-treated versus TGFβ treated THP-1 cells, mock versus MCEMP1 overexpressing cells treated with TGFβ and sh control versus shMCEMP1 treated with TGFβ. Differentially expressed genes (DEG) with P<0.05 (FDR adjusted) were selected for downstream analyses.

### Active RHO pull-down assay

THP-1 cells were stably transduced with control or MCEMP1-targeting shRNA using lentivirus and cultured in 75 cm ^2^ flasks to reach at least 10 million cells per flask. Cells were further cultured with or without TGFβ (5ng/ml) for 24 hours, and pelleted by centrifugation (100 x g, 5 min). After resuspension in ice-cold TBS, cells were centrifuged, lysed and the supernatant was collected to measure protein concentration. For each sample, 1mg of protein was used to pull down active RHO (RHO-GTP) using the Active Rho Pull-Down and Detection kit from Thermofisher by following the manufacturer’s instructions. A volume of each cell lysate was used for detection of total Rho, MCEMP1 and GAPDH by western blot.

### Statistical analysis

Statistics were performed using GraphPad Prism 10. The two-tailed Student’s t test was used to compare two conditions. For multiple comparisons, we used either the one-way or two-way ANOVA with Dunnett’s, Tukey’s or Sidak multiple comparison test as appropriate for each comparison. Error bars represent standard deviation (SD). Graphic definition of statistical significance: ****p<0.0001, ***p<0.001, **p<0.01, *p<0.05.

### Statistical analysis of bulk RNA sequencing data

Differential gene expression analysis was performed using Deseq2 package(12). Three groups were analyzed: non-treated versus TGFβ treated cells, MCEMP1 mock versus MCEMP1 overexpressing cells treated with TGFβ and sh control versus shMCEMP1 treated with TGFβ. Differentially expressed genes (DEG) with P<0.05 (FDR adjusted) were selected for downstream analyses. Gene Ontology (GO) analysis was performed on Bulk RNA sequencing. Statistical significance for GO analysis was defined as Bonferroni corrected P<0.05.

## Results

### MCEMP1 is highly expressed in circulating classical monocytes and alveolar macrophages in IPF when compared to controls

To understand the cellular source of MCEMP1 expression in IPF and whether its expression was different from age and gender matched healthy controls, we analyzed scRNA-seq of PBMC from IPF patients and controls (9). Table 1 summarizes the demographic differences of the studied subjects. Our analysis indicated that classical monocytes are the main cellular source of MCEMP1 in PBMC (Figure 1A). While MCEMP1 expression is higher in classical monocytes, it is also detected (albeit modestly) in intermediate monocytes, dendritic cells, and thrombocytes, however, MCEMP1 is not expressed in non-classical monocytes and in lymphocytes in IPF (Figure 1B). In circulating classical monocytes, MCEMP1 expression was significantly higher in patients with IPF compared to control subjects (Figure 1C).

**Figure 1.**
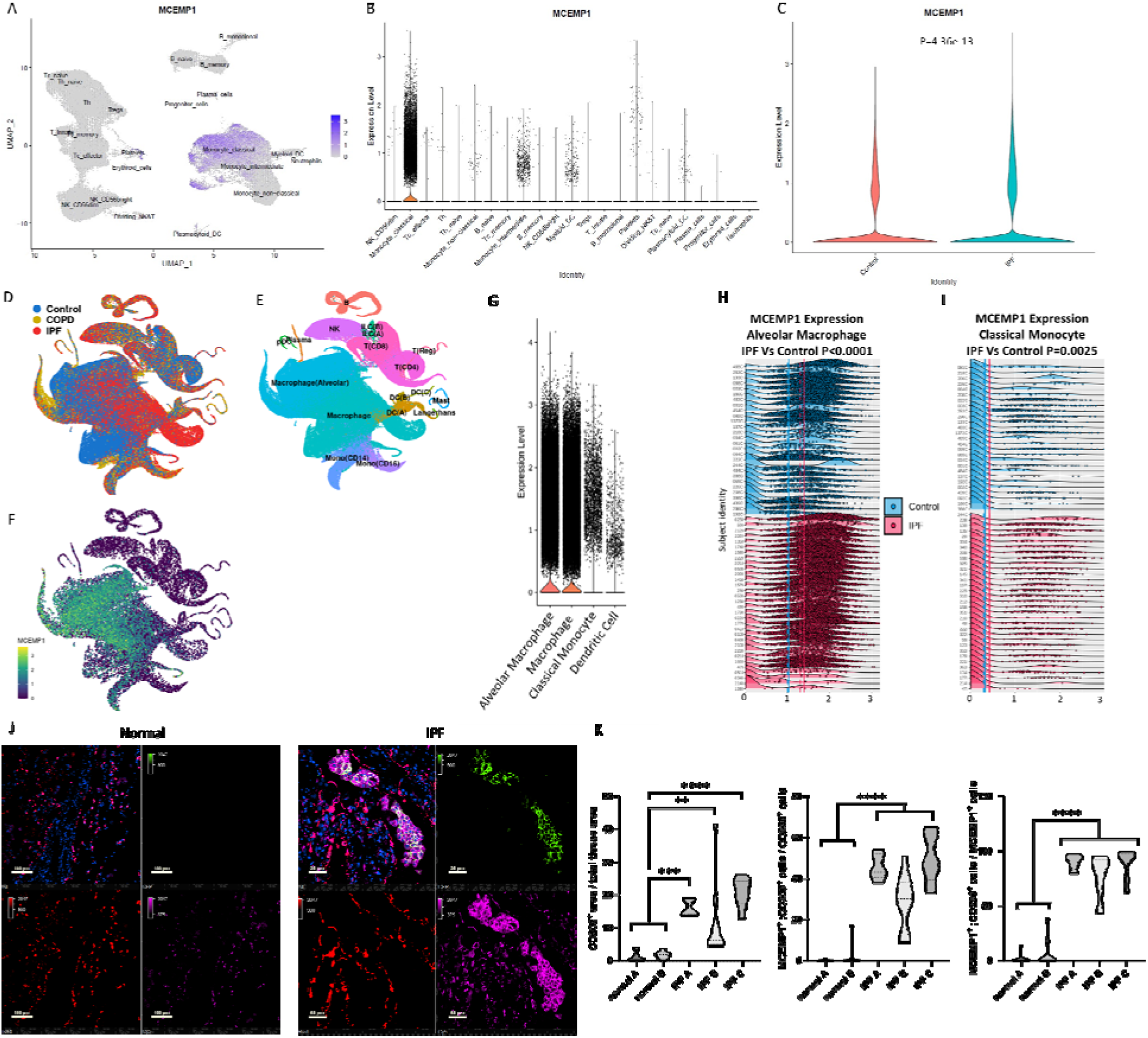
MCEMP1 expression is increased in circulating classical monocytes and alveolar macrophages in IPF patients compared to controls. UMAP representation of 149,564 cells parceled into 23 cell types. All expected cell types are included. MCEMP1 expression is colored purple and color scale is adjacent to the UMAP. MCEMP1 expression is highest in classical monocytes in IPF (A). Other cell types expressing MCEMP1 to a lesser extent include intermediate monocytes, platelets, myeloid-derived and plasmacytoid dendritic cells (B). Violin plots demonstrate that MCEMP1 expression in classical monocytes is significantly increased in IPF (green) versus controls (red) (C). Wilcoxon rank sum test, P<0.0001. UMAP projections of RNA sequencing of 312,928 single cells from Control, Chronic Obstructive Pulmonary Disease (COPD) and Idiopathic Pulmonary Fibrosis (IPF) lung tissues (E). Density plots show cell population density based on cellular markers (F) and MCEMP1 expression (G). High expression of MCEMP1 is detected in Alveolar Macrophages, Macrophages, CD14 ^+^ Monocytes and Dendritic Cells (H). MCEMP1 expression is significantly higher in Alveolar Macrophages and in Classical Monocytes in IPF patients when compared to controls (I). Y axis (subject identity), X axis (expression levels). Immunofluorescence staining demonstrates absence of MCEMP1 staining in normal lungs and its presence and co-localization in alveolar macrophages in IPF (J). The difference between cells co-expressing CD206 and MCEMP1 was significantly higher in IPF compared to controls (K). DAPI: Blue, MCEMP1: Green, HT2-280: Red and CD206: Purple. ****p<0.0001, ***p<0.001, **p<0.01

**Table 1.** Baseline characteristics of IPF patients and control subjects in the PBMC scRNA-se q dataset.

|  | IPF | Controls | P value |
| --- | --- | --- | --- |
| N | 25 | 13 |  |
| Age mean (SD), years | 70.3 (6.4) | 71.0 (4.2) | 0.70 |
| Male sex | 80% | 77% | 0.83 |
| Caucasian race | 96% | 100% | 0.46 |
| Current/former smoker | 72% | 69% | 0.86 |
| FVC, % predicted | 73.48 | NA | NA |
| DLCO, % predicted | 41.12 | NA | NA |

To determine the cellular source of MCEMP1 in IPF lung tissues we analyzed scRNA-seq cells of the distal lung parenchyma from 32 IPF lungs, 18 chronic obstructive pulmonary disease (COPD) and 28 control donor lungs (10). We identified that MCEMP1 is primarily expressed in lung tissue in alveolar macrophages, macrophages, classical monocytes (CD14 ^+^) and dendritic cells (Figures 1D-G). MCEMP1 expression was significantly higher in alveolar macrophages (Figure 1H) and classical monocytes (Figure 1I) in IPF when compared to control lungs. To validate our findings, we performed immunofluorescence staining and co-localization of MCEMP1 with CD206 in IPF lung tissue and control lungs. MCEMP1 staining was essentially absent in control lungs while it was detected and co-localized with CD206 in alveolar macrophages in IPF (Figure 1J). The differences in the total number of CD206 ^+^MCEMP1^+^ cells between IPF and controls by immunofluorescence staining was statistically significant (Figure 1K).

### MCEMP1 is a TGFβ inducible gene in monocytes

Once we confirmed classical monocytes as one of the predominant MCEMP1-expressing cells in IPF, we used the monocytic cell line THP-1 as an *in vitro* model to elucidate the molecular mechanisms regulating the expression of this gene. THP-1 cells were treated with several compounds known to promote inflammation and/or fibrosis: Tumor Necrosis Factor -α (TNFα, 10ng/ml), Fibroblast Growth Factor-2 (FGF2, 100ng/ml), TGFβ (5ng/ml) or lipopolysaccharide (LPS, 50ng/ml). We analyzed MCEMP1 levels by RT-qPCR and western blot 24 hours post-treatment and we found that TGFβ increased MCEMP1 both at the mRNA and at the protein levels, while TNFα, FGF2 and LPS did not have any effect (Figures 2A, B). THP-1 cells were also stimulated with profibrotic interleukins-4 and -13 and proinflammatory interleukin-10 (10ng/ml), but no effect on MCEMP1 expression was seen (Figure 2C). MCEMP1 overexpression happened as early as 12 hours. Phospho-SMAD2 and total SMAD2 protein levels were analyzed by western blot as a positive control for TGFβ efficiency and levels were noted to be increased at 12 and 24 hours (Figure 2D). The observation that TGFβ induces MCEMP1 expression was confirmed by live cell imaging of THP-1 cells immunostained with an MCEMP1 antibody and with a CD14 antibody as a control (Figure 2E).

**Figure 2.**
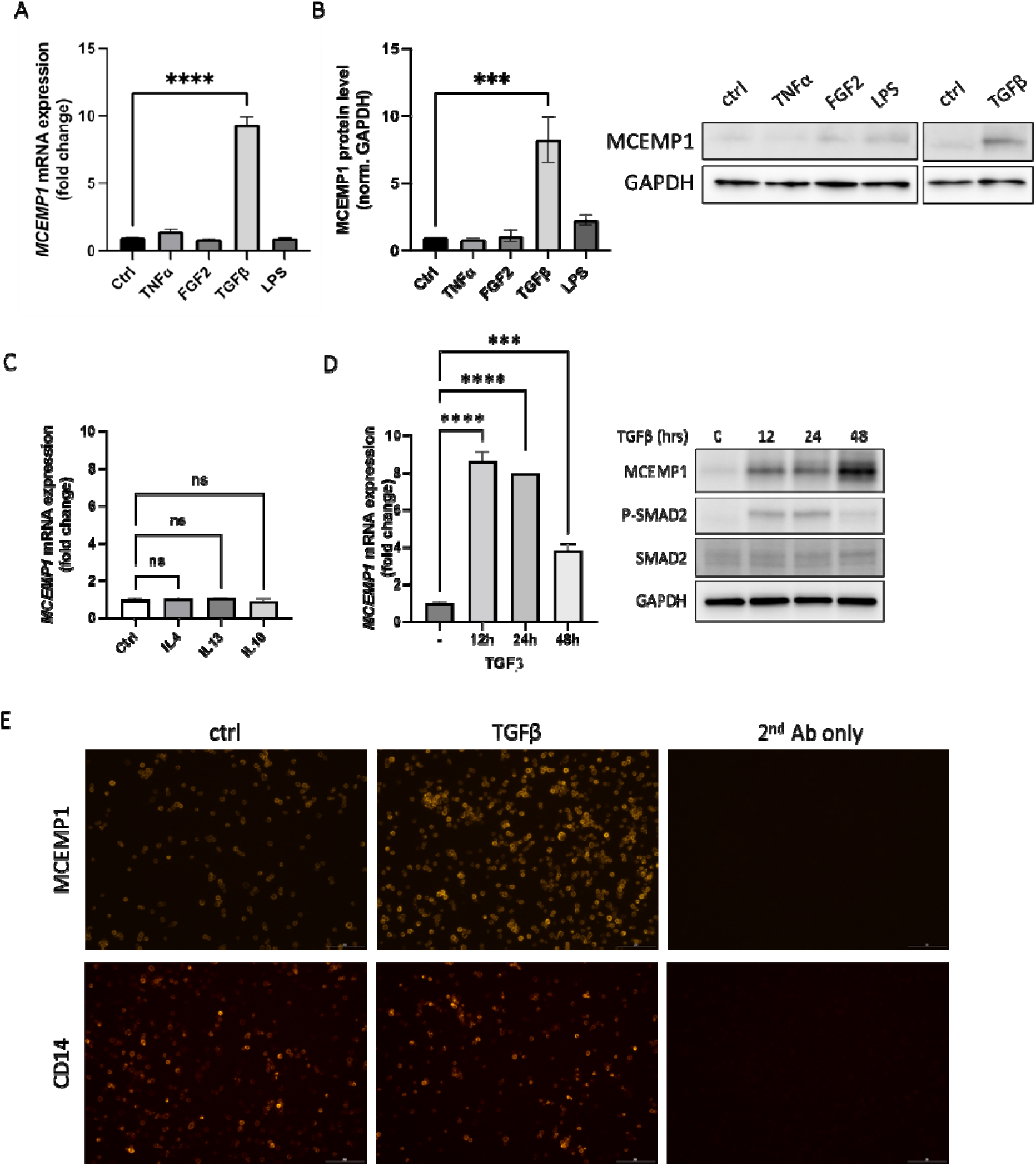
MCEMP1 is a TGFβ inducible gene in monocytes. MCEMP1 was found to be upregulated by TGFβ at the gene expression and protein levels (A). Stimulating THP-1 cells with Fibroblast growth factor 2 (FGF2), Tumor necrosis factor alpha (TNF-α) and lipopolysaccharide (LPS) did not significantly change MCEMP1 RNA and protein levels (B). Interleukins 4, 13 and 10 did not affect MCEMP1 expression levels. The observation that TGFβ increased MCEMP1 was confirmed and by live cell immunostaining (E). One-way ANOVA, ****p<0.0001, ***p<0.001. Scale bar 200μm.

### TGFβ-mediated MCEMP1 upregulation results from the cooperation of SMAD3 and SP1 via concomitant binding to SMAD3/SP1 cis-regulatory elements within the MCEMP1 promoter region

We used the NIH genetic sequence database GenBank to extract and analyze the sequence of a 1.5kbp promoter region upstream of the human *MCEMP1* gene ATG start codon to identify SMAD-binding element (SBE) and CAGA boxes (Figure 3A). Our analysis was set up with a minimum sequence homology of 0.7 and a core sequence homology of 1, which resulted in the identification of three putative SBE/CAGA sites at locations -1,158/-1,152 (TCT**AGAC**AGA), -937/-931 (**AGAC**AGA) and -325/-314 (**AGAC**AC**AGAC**ACACACAG**AGAC**). The proximal part of *MCEMP1* promoter is a GC-rich region with a GC-box at position -79/-73 (GGGCGGG; sequence homology 1, core sequence homology 1) (Figure 3A).

**Figure 3.**
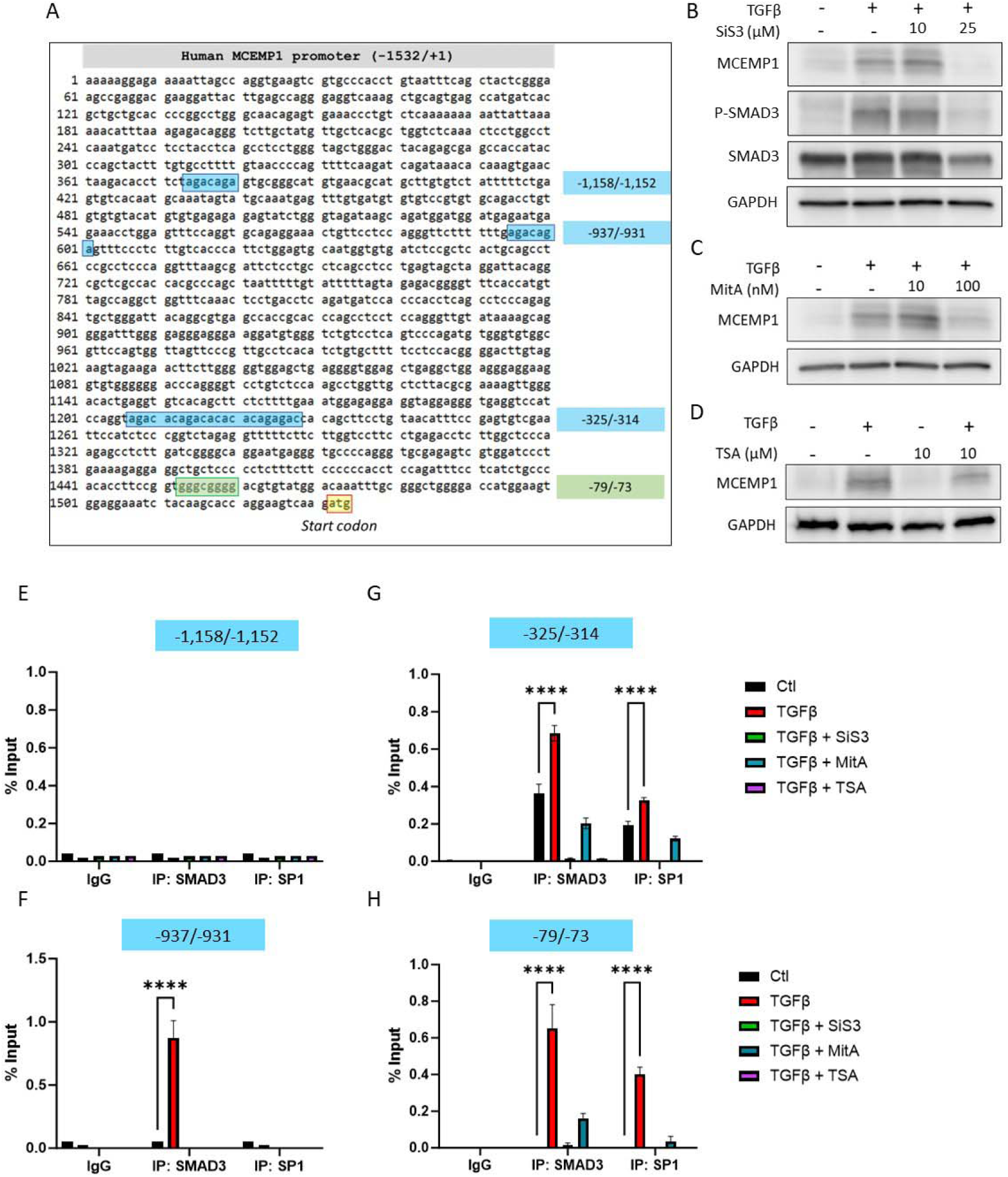
TGFβ-mediated MCEMP1 upregulation results from the cooperation of SMAD3 and SP1 via concomitant binding to SMAD3/SP1 *cis*-regulatory elements within the *MCEMP1* promoter region. Analysis of the sequence of a 1.5kbp promoter region upstream of the human *MCEMP1* gene ATG start codon resulted in the identification of three putative SBE/CAGA sites at locations -1,158/-1,152 (TCT **AGAC**AGA), -937/-931 (**AGAC**AGA) and -325/-314 (**AGAC**AC**AGAC**ACACACAG**AGAC**) and a GC box at position -79/73 (GGGCGGG) (A). TGFβ failed to upregulate MCEMP1 expression in the presence of the SMAD3 inhibitor SiS3 (B) or SP1 inhibitor mithramycin A (MitA) (C). Similarly, THP-1 treatment with trichostatin A (TSA), also blocked MCEMP1 upregulation following TGFβ stimulation (D). Chromatin immunoprecipitation followed by qPCR shows that neither SMAD3 nor SP1 can bind to the -1,158/-1,152 SBE/CAGA site (E). SMAD3 binds to the -937/-931 SBE/CAGA site, but not SP1 (F). Concomitant binding of SMAD3 and SP1 is observed at locations -325/-314 and -79/-73 following THP-1 stimulation with TGFβ. This effect is inhibited in the presence of SiS3 or MitA (G, H).

To determine if SMAD3 and/or SP1 are transcriptional effectors in the induction of MCEMP1 expression by TGFβ, THP-1 cells were treated either with SMAD3 inhibitor SiS3 or SP1 inhibitor mithramycin A (MitA), prior to TGFβ stimulation. Twenty-four hours later, we observed that TGFβ failed to increase MCEMP1 expression in the presence of either inhibitor (Figure 3B, C). This suggests that TGFβ requires both SMAD3 and SP1 to induce MCEMP1 expression. Similarly, THP-1 treatment with trichostatin A (TSA), a histone deacetylase (HDAC) inhibitor known to block SP1 function, also blocked MCEMP1 upregulation following TGFβ stimulation, confirming the pivotal role of SP1 in this process (Figure 3D)(13).

To assess SMAD3 and SP1 binding to the *MCEMP1* promoter in THP-1 cells, we performed ChIP to scan the regions spanning the *cis*-regulatory elements identified *in silico.* qPCR amplification following SMAD3 and SP1 immunoprecipitation revealed that SMAD3, but not SP1, is recruited at site -937/-931 following TGFβ stimulation (Figure 3F). This effect was abrogated in the presence of SiS3. Both SMAD3 and SP1 were found to bind to site -325/-314 and to the GC-box (-79/-73), but not in the presence of SiS3, MitA and TSA (Figure 3G, 3H). Neither SMAD3 nor SP1 binding was detected on the predicted -1,158/-1,152 SBE/CAGA site (Figure 3E). Taken together, our results demonstrate that TGFβ-mediated MCEMP1 overexpression results from the cooperation of SMAD3 and SP1 via concomitant binding to SMAD3/SP1 *cis*-regulatory elements within the MCEMP1 promoter.

### Proximity ligation assay identifies the interaction of SMAD3 and SP1 *in situ*, which is enhanced by TGF-β treatment

To visualize SMAD3 and SP1 interaction in THP-1 cells, we performed a proximity ligation assay (PLA) in THP-1 cells cultured and treated with TGFβ in the presence or absence of SiS3 or mithramycin A. We used primary antibodies targeting each protein first, and secondary antibodies labeled with oligonucleotide probes. In our experiment, we observed an important increase of fluorescent spots following TGFβ treatment compared to control, particularly in nuclei where both transcription factors translocate to regulate gene expression (Figures 4A, B, E). Both SiS3 and MitA drastically reduced the formation of SMAD3-SP1 complexes in THP-1 nuclei, demonstrating the specificity of the PLA signal (Figures 4C, D, E).

**Figure 4.**
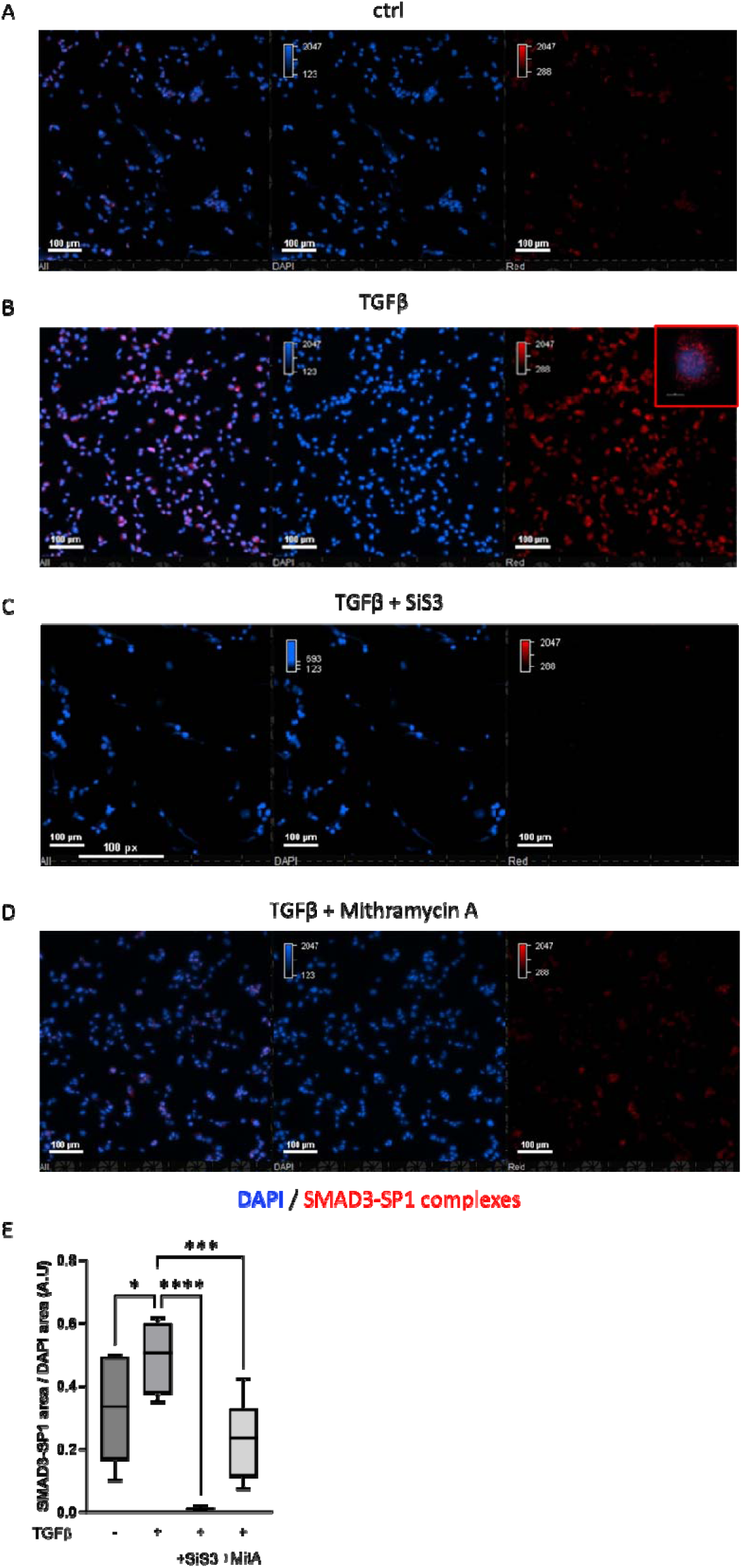
Proximity ligation assay identifies the interaction of SMAD3 and SP1 *in situ*, which is enhanced by TGF-β treatment. TGFβ treatment resulted in an important increase of fluorescent spots following TGFβ treatment compared to control, particularly in nuclei where both transcription factors translocate to regulate gene expression (A, B, E). Both SiS3 and MitA drastically reduced the formation of SMAD3-SP1 complexes in THP-1 nuclei, demonstrating the specificity of the PLA signal (C, D, E). DAPI (Blue), SMAD3-SP1 complexes (Red). ****p<0.0001, ***p<0.001, *p<0.05.

### MCEMP1 regulates TGFβ-mediated monocyte chemotaxis, adhesion, and migration

Once we determined that MCEMP1 expression was detected in classical monocytes and alveolar macrophages in IPF, and regulated by TGFβ signaling, we then hypothesized that MCEMP1 could participate in the transition of circulating monocytes to monocyte-derived alveolar macrophages by controlling cell chemotaxis and/or migration in a TGFβ rich environment. To test our hypothesis, we performed a cell migration/chemotaxis assay in in THP-1 cells with either increased or decreased expression of MCEMP1 using transwells and CCL2 (MCP-1) as a chemoattractant. The efficiency of transfection and transduction for MCEMP1 knockdown and knock-in was confirmed by western blot (supplementary Figure S1A-D). As shown on Figure 5A, THP-1 cells did not cross the transwell membrane in the absence of CCL2. In the presence of CCL2, we observed a dramatic increase in transwell migration upon TGFβ stimulation. This effect was reduced by approximately 75% following *MCEMP1* knockdown. To determine whether MCEMP1 overexpression could increase THP-1 chemotaxis and migration in the absence of TGFβ, we stably transduced THP-1 cells with mock- and *MCEMP1*-lentiviral vectors to induce MCEMP1 expression. We then performed a chemotaxis/migration assay using the same system as in Figure 5A and tested cells from two different transductions as biological replicates, both for mock- and MCEMP1-lentivirus. We observed that MCEMP1-overexpressing cells from both transductions migrated more than mock control cells, which indicates that MCEMP1 enhances THP-1 chemotaxis (Figure 5B).

**Figure 5.**
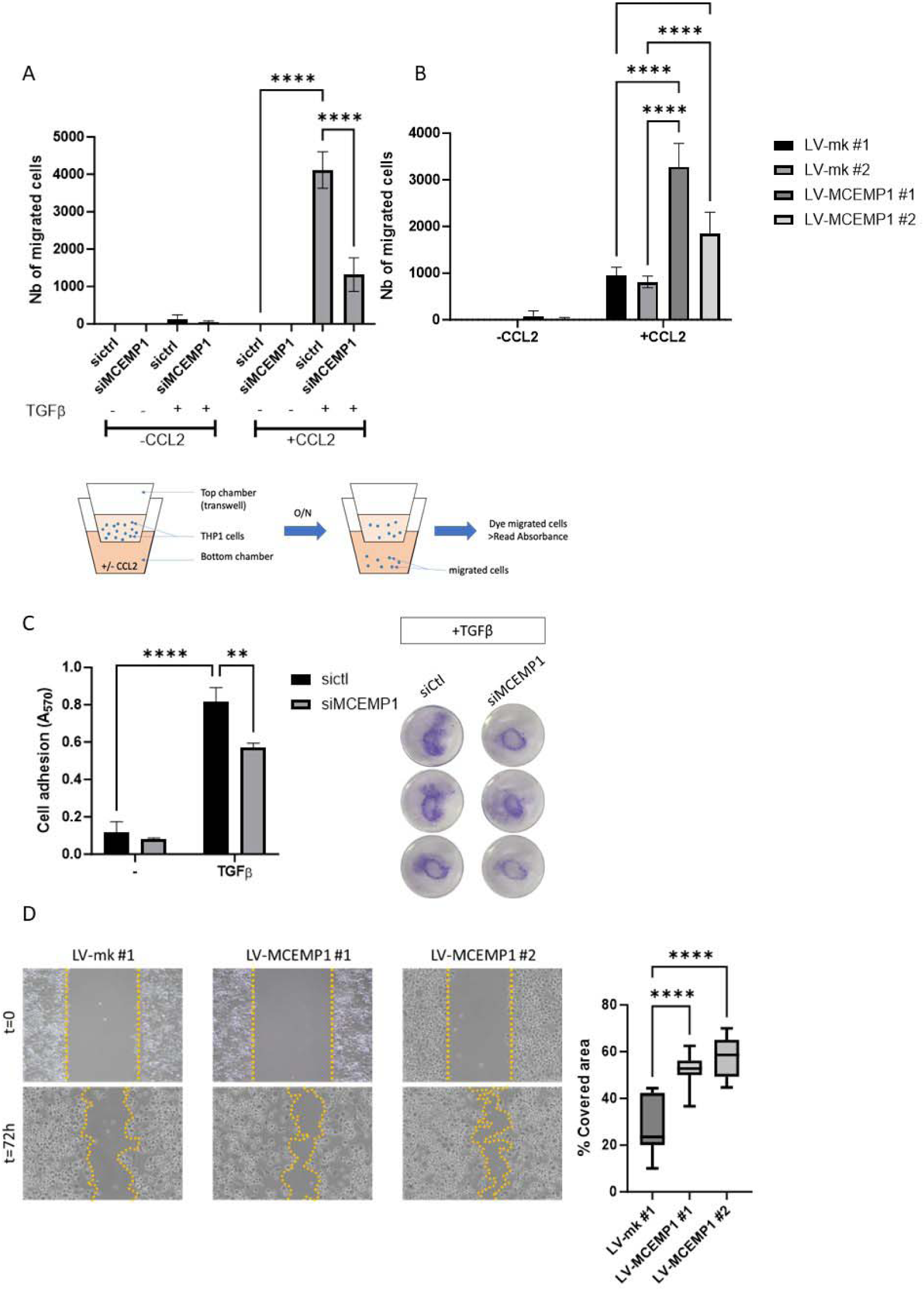
MCEMP1 regulates TGFβ-mediated monocyte chemotaxis, adhesion, and migration. THP1 cells treated with TGFβ and transfected with *MCEMP1*-targeting siRNA migrate significantly less after CCL2 stimulation (A). THP1 cells stably transduced with MCEMP1-encoding lentiviral vectors (LV-MCEMP1) showed increased migration when compared to cells transduced with an empty vector. Two different clones were tested (B). While TGFβ treatment potently increased THP-1 attachment to the cell culture dish, this effect was partially impaired by *MCEMP1* knockdown (C). Scratch assay results show that MCEMP1 overexpression increases macrophage migration capacity compared to control cells (D). Two-way ANOVA, ****p<0.0001, ***p<0.001, **p<0.01

Since TGFβ promotes monocyte adhesion, we tested whether MCEMP1 could be involved in this process by performing a cell adhesion assay using control- or MCEMP1-siRNA transfected THP-1 cells, stimulated or not with TGFβ (14). As expected, TGFβ treatment potently increased THP-1 attachment to the cell culture dish. However, this effect was partially impaired by MCEMP1 knockdown, which suggests a role for MCEMP1 in TGFβ-mediated monocyte adhesion (Figure 5C). The overexpression of MCEMP1 without addition of TGFβ failed to induce THP-1 adhesion (data not shown). These observations demonstrate that MCEMP1 partially controls TGFβ-mediated monocyte adhesion in THP-1 cells.

Next, we aimed to check if MCEMP1 could control THP-1 migration following differentiation into macrophages. We seeded mock- and MCEMP1-overexpressing THP-1 in six-well culture plates and treated them with PMA to induce cell differentiation. The results of a scratch assay showed that MCEMP1 overexpression increases macrophage migration capacity compared to control cells. Altogether, our experiments provide evidence that MCEMP1 regulates monocyte and macrophage chemotaxis, adhesion, and migration, upon exposure to TGFβ.

### MCEMP1 mediates RHO-GTPase activity following TGFβ stimulation

To study the mechanisms by which MCEMP1 participates in monocyte motility, we performed bulk RNA sequencing in three groups of THP1 cells: non-treated versus TGFβ treated cells, MCEMP1 mock versus MCEMP1 overexpressing cells treated with TGFβ (knock-in MCEMP1) and sh-control versus shMCEMP1 cells treated with TGFβ (knockdown MCEMP1). Differentially expressed genes (DEG) with P<0.05 (FDR adjusted) were selected for downstream analyses. We identified N=400 DEG that increase after TGFβ stimulation, further increased in MCEMP1 overexpressing cells treated with TGFβ and decreased in shMCEMP1 cells treated with TGFβ (Figure 6A). We also identified N=196 genes with the opposite pattern of expression (Figure 6B). For enrichment analyses, we focused on the former groups of DEG genes (Figure 6A). GO analysis of biological processes demonstrated enrichment for positive regulation of GTPase activity and signal transduction (Table 2).

**Figure 6.**
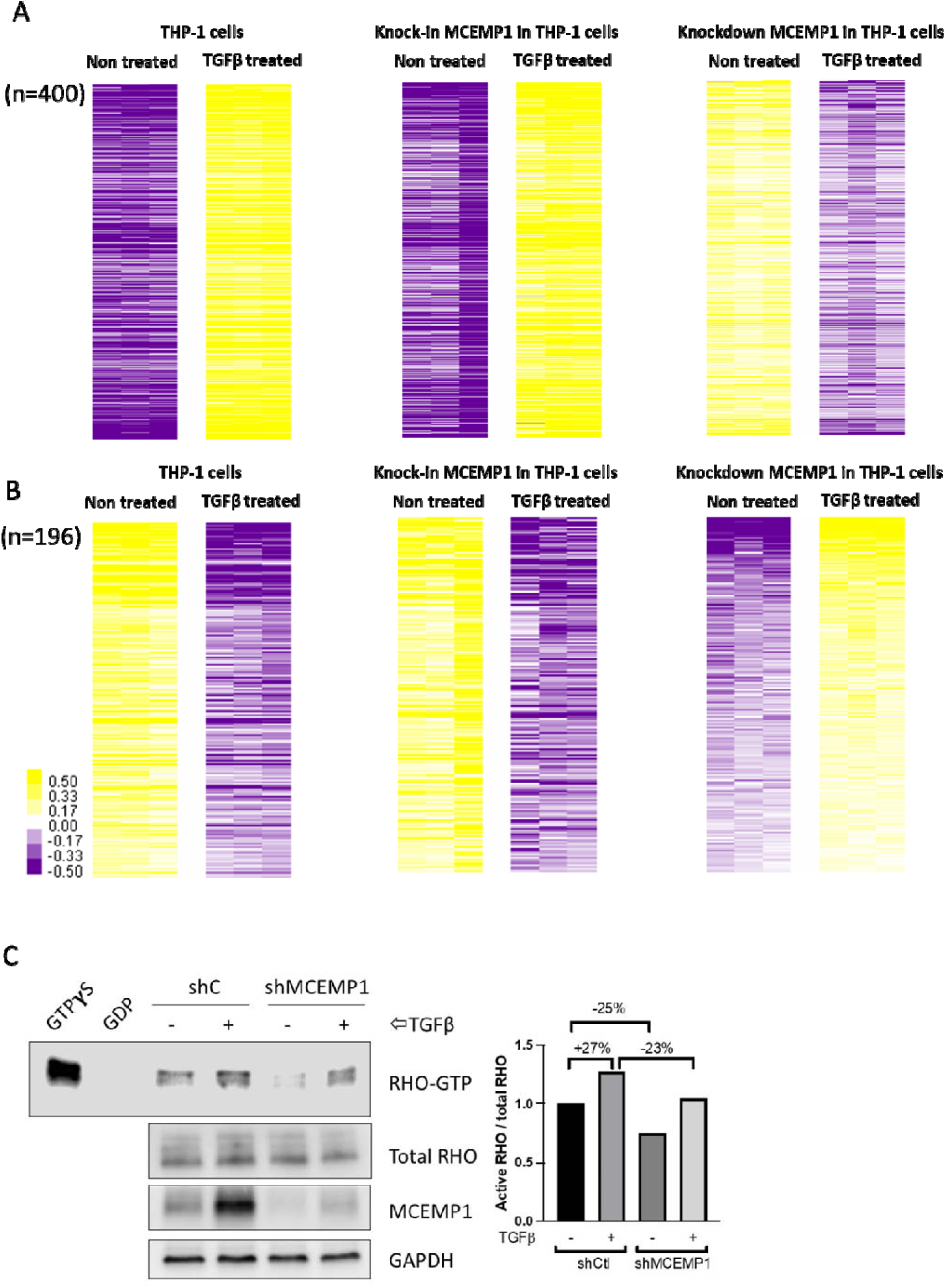
MCEMP1 mediates RHO-GTPase activity following TGFβ stimulation. . Heatmap depicts Differentially Expressed Genes (DEG) from bulk RNA sequencing of three groups of THP1 cells. N=400 DEG were found to increase after TGFβ stimulation, further increased in MCEMP1 overexpressing cells treated with TGFβ and decreased in shMCEMP1 cells treated with TGFβ (A). N=196 genes followed the opposite pattern of expression (B). Color scale is shown adjacent to heat map in log _2_ scale; generally, yellow denotes increase over the geometric mean of samples, and purple, decrease. Differential gene expression analysis based on the negative binomial distribution. FDR < 5%. THP1 cells transfected with *MCEMP1*-targeting shRNA (shM) and treated with TGFβ have a 23% reduction in active RHO-GTPase activity compared to sh control cells treated with TGFβ (C).

**Table 2.** GO annotation analysis of N=400 differentially expressed genes responsive to TGF β and regulated by MCEMP1 in THP-1 cells show enrichment for positive regulation of GTPas e activity and signal transduction. Bonferroni corrected P<0.05.

| GO biological process | Genes expected | Genes identified | Fold change | P-value |
| --- | --- | --- | --- | --- |
| Establishment of epithelial cell polarity (GO:0090162) | 35 | 8 | 11.71 | 1.26E-02 |
| Transforming growth factor beta receptor signaling pathway (GO:0007179) | 96 | 13 | 6.94 | 1.58E-03 |
| Cellular response to transforming growth factor beta stimulus (GO:0071560) | 155 | 15 | 4.96 | 9.57E-03 |
| Response to transforming growth factor beta (GO:0071559) | 162 | 15 | 4.74 | 1.59E-02 |
| Regulation of small GTPase mediated signal transduction (GO:0051056) | 304 | 28 | 4.72 | 5.71E-07 |
| bone development (GO:0060348) | 220 | 19 | 4.42 | 1.82E-03 |
| Positive regulation of GTPase activity (GO:0043547) | 278 | 22 | 4.05 | 8.26E-04 |
| Response to hypoxia (GO:0001666) | 274 | 20 | 3.74 | 1.04E-02 |
| Endocytosis (GO:0006897) | 534 | 38 | 3.64 | 2.78E-07 |
| Regulation of GTPase activity (GO:0043087) | 368 | 26 | 3.62 | 4.69E-04 |

RHO GTPases such as RHO, RAC, and CDC42 have been shown to play an essential role in the regulation of monocyte/macrophage chemotaxis and migration in response to chemokines such as CCL2 or colony-stimulating factor-1 (CSF-1)(15). Therefore, we monitored RHO activation following TGFβ stimulation in THP-1 stably transduced with control- or MCEMP1-shRNA. We found that TGFβ enhances RHO GTPase activity, however this effect was attenuated following MCEMP1 knockdown (Figure 6C). Taken together, these results support the role of MCEMP1 in TGFβ-mediated motility of monocytes through the modulation of RHO activity.

## Discussion

Our study provides both human data and strong *in-vitro* evidence for the role of MCEMP1 in the pathogenesis of pulmonary fibrosis. We showed through single-cell RNA sequencing that MCEMP1 is expressed primarily in classical monocytes and in alveolar macrophages in IPF. Thus, we used a monocytic cell line (THP-1 cells) for our mechanistic studies and demonstrated that MCEMP1 is a transcriptional target of TGFβ, a key mediator of pulmonary fibrosis. TGFβ transduces signals through canonical and non-canonical pathways (16–18). The well-characterized canonical TGFβ signaling pathway uses SMAD2 and/or SMAD3 along with SMAD4 to transfer signals, and SMAD3 and SMAD4 are able to directly bind to DNA through two different consensus sequences: GTCTAGAC, called SMAD binding element or SBE; and AG(C/A)CAGACAC, called CAGA box. Both sequences contain the core motif **<u>AGAC</u>**, which represents the optimal binding sequence for SMAD3 and SMAD4 (19, 20). Here we show that TGFβ signaling regulates MCEMP1 expression in a non-canonical fashion by enhancing SMAD3 and SP1 interaction and concomitant binding to SMAD3/SP1 *cis*-regulatory elements in positions -325/-314 and -79/-73 within the *MCEMP1* promoter. These results corroborate previous studies describing SMAD3-SP1 transcriptional complexes (21, 22). Additionally, our study provides evidence that MCEMP1 enhances TGFβ-mediated monocyte adhesion, chemotaxis, and migration by modulating the effect of TGFβ in RHO activity.

Once we determined that MCEMP1 expression is regulated by TGFβ-SMAD3-SP1 and mainly detected in monocytes, we sought to understand the function of the protein. Monocytes are cells with migratory capacity, being in the circulation, but they also leave this compartment and migrate to peripheral tissues where they differentiate into macrophages (23). We thus speculated and showed that MCEMP1 is involved in monocyte trafficking by controlling chemotaxis and/or migration. Given that TGFβ promotes monocyte migration and adhesion as well macrophage migration through Rho GTPases, we tested whether MCEMP1 could be involved in this process (15, 24–27). Our results provide evidence that increased MCEMP1 expression following monocyte and macrophage exposure to TGFβ enhances cell chemotaxis, adhesion, and migration by regulating the effect of TGFβ in RHO GTPase signaling. Finally, we identified the presence of MCEMP1 in lung tissue monocytes and macrophages, particularly in alveolar macrophages in IPF, suggesting that these cells could be monocyte derived and that MCEMP1 may participate in the migration and transition of monocytes to monocyte-derived alveolar macrophages. Further in vivo studies will be required to demonstrate that alveolar macrophages in IPF expressing MCEMP1 are monocyte derived.

Our findings showing a role of MCEMP1 in THP-1 cell chemotaxis and migration is in line with mechanistic data of MCEMP1 function in the context of other diseases (28, 29). In particular, it has been demonstrated that MCEMP1 may affect the proliferation, migration, and invasion of gastric cancer cells through regulating epithelial to mesenchymal transition (29). It is also known that small GTPases of the Rho/Rac family participate in TGF-β-induced EMT and cell motility in cancer(27), an effect that based on our results seems to be mediated in monocytes and macrophages in IPF, at least in part by MCEMP1. In addition to pulmonary fibrosis, a recent study demonstrated that MCEMP1 deficiency blunted SCF-induced peritoneal mast cell proliferation *in vitro* and lung mast cell expansion *in vivo* . *Mcemp1*-deficient mice had decreased airway inflammation and lung impairment in mouse models of chronic asthma (30). These findings highlight that MCEMP1 might be an emerging key mediator for multiple diseases. Additionally, elevated MCEMP1 has been recently reported as a negative prognostic marker in COVID-19 as well as other diseases such as stroke, thus is possible that MCEMP1 participates in fibrosis post injury (31–33).

Our study has some limitations. First, we did not use primary human monocytes for our mechanistic experiments, but a monocytic cell-line (THP-1 cells). Primary monocytes are short lived and difficult to transfect or transduce. We used THP-1 cells because MCEMP1 was first described in THP-1 cells (8). THP-1 cells exhibit a fast growth rate, are homogenous in cell culture and can differentiate. Also, THP-1 cells are a widely used and validated model to study monocyte biology (34) . Second, we did not validate our results *in vivo*; yet we provide strong *in vitro* evidence for the role of MCEMP1 in pulmonary fibrosis.

Taken together, our study has several important attributes. Our data demonstrates that MCEMP1, a gene of a 52-gene, high-risk profile predictive of IPF mortality when upregulated, is highly expressed in classical monocytes in PBMC and alveolar macrophages in IPF lung tissues when compared to control lungs. This is also the first study providing mechanistic insights for the role of MCEMP1 in pulmonary fibrosis and one of the few studies identifying the mechanistic role of MCEMP1 in general. Our study not only identified that MCEMP1 expression is regulated by the TGFβ-SMAD3-SP1 axis but also that MCEMP1 seems to be involved in the crosstalk between TGFβ and RHO-GTPase in monocyte chemotaxis and macrophage migration, a mechanism that has not been previously reported. The findings noted above imply a potentially important mechanistic role of MCEMP1 for pulmonary fibrosis development and progression. Mcemp1 knock-in and knockout mouse models will be required in future studies to further validate our *in vitro* findings.

## Declarations

### Ethics approval

Protocol, data collection and analysis were approved by the Institutional Review Board and the Local Ethics Committee (BullsIRB protocol: Pro00032158).

### Consent for publication

Consent for publication by all authors.

### Availability of data and materials

Data available upon request.

### Contributors - Author contribution statement

**Carole Y. Perrot:** conceptualisation, investigation, methodology, data curation, visualisation, resources, writing – original draft

**Theodoros Karampitsakos** : conceptualisation, investigation, methodology, data curation, visualisation, writing – original draft

**Avraham Unterman:** investigation, methodology, data curation, writing- review& editing

**Taylor Adams:** investigation, methodology, data curation, writing- review& editing

**Krystin Marlin:** investigation, methodology, data curation, writing- review& editing

**Alyssa Arsenault:** investigation, methodology, data curation, writing- review& editing

**Amy Zhao:** investigation, methodology, data curation, writing- review& editing

**Naftali Kaminski:** conceptualisation, investigation, methodology, data curation, supervision, funding acquisition, project administration, writing- review& editing

**Gundars Katlaps:** investigation, methodology, data curation, writing- review& editing

**Kapilkumar Patel:** investigation, methodology, data curation, writing- review& editing

**Debabrata Bandyopadhyay:** investigation, methodology, data curation, writing- review& editing

**Jose D. Herazo-Maya:** conceptualisation, investigation, methodology, data curation, visualisation, project administration, supervision, resources, funding acquisition, project administration, writing – original draft

All authors had substantial contribution to the following: (1) the conception and design of the study, or acquisition of data, or analysis and interpretation of data, (2) drafting the article or revising it critically for important intellectual content, (3) final approval of the version to be submitted. Carole Y. Perrot and Theodoros Karampitsakos contributed equally to this work.

## Acknowledgments

none to declare.

