## Supplementary data for "Mast-Cell Expressed Membrane Protein-1 (MCEMP1) is expressed in classical monocytes and alveolar macrophages in Idiopathic Pulmonary Fibrosis and regulates cell chemotaxis, adhesion, and migration in a TGFβ dependent manner"

*Equal contribution

Correspondence to:

University of South Florida

**Supplementary Figure**

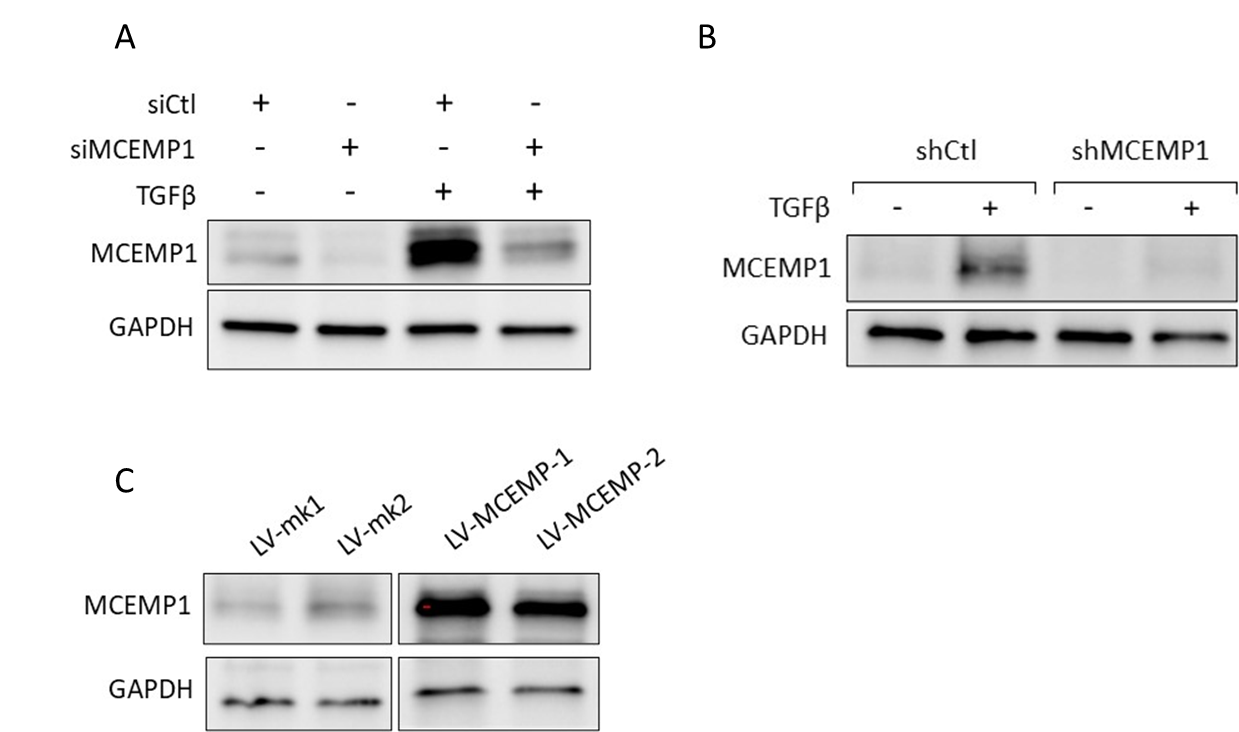

**Supplementary Figure S1**

*MCEMP1* silencing efficiency using siRNA was verified by western blot (A). MCEMP1 stable knockdown in THP-1 using shMCEMP1 lentivirus was confirmed by western blot. TGFβ failed to upregulate MCEMP1 in shMCEMP1 cells (B). MCEMP1 overexpression using lentiviral vectors was confirmed by western blot. Two clones were tested (LV-MCEMP-1 and LV-MCEMP1-2) (C).

**Supplementary Table**

| **Primer sets for qPCR** | | |
| --- | --- | --- |
| Primers | sequence | |
| MCEMP1-Fw | 5' | GTACATGTGGGTGTCGCCAT |
| MCEMP1-Rv | 5' | TGGCATAACTCGGCACACTC |
| RPS18-Fw | 5' | CGATGGGCGGCGGAAAATA |
| RPS18-Rv | 5' | CTGCTTTCCTCAACACCACA |
| RPL37A-Fw | 5' | ATCTGGCACTGTGGTTCCTG |
| RPL37A-Rv | 5' | GACTTTACCGTGACAGCGGA |
| **Primer sets for ChIP** | | |
| Primers | sequence | |
| CAGA1-Fw (-1158/-1152) | 5' | GTAACCCCAGTTTTCAAGATCAG |
| CAGA1-Rv (-1158/-1152) | 5' | GAAAAATAGACACAAGCATGCG |
| CAGA2-Fw (-937/-931) | 5' | GCAGATGGATGGATGAGAATGAG |
| CAGA2-Rv (-937/-931) | 5' | GAGATCACACCATTGCACTCCA |
| CAGA3-Fw (-325/-314) | 5' | GAGGTGTCACAGCTTCTCTTTTG |
| CAGA3-Rv (-325/-314) | 5' | ATGGAATTCGACACTCGGAAATG |
| GC-BOX-Fw (-79/-73) | 5' | TGCGAGAGTCGTGGATCCCTGA |
| GC-BOX-Rv (-79/-73) | 5' | GTGCTTGTAGATTTCCTCCACTTC |
| **siRNA and shRNA sequences** | | |
| siMCEMP1 sense | 5' | GAAUGUCUCAAACUCCGUAtt |
| siMCEMP1 antisense | 5' | UACGGAGUUUGAGACAUUCca |
| shMCEMP1 | 5' | ATCTACAAGCACCAGGAAGTCAAGATGCA |

**Supplementary Table S1**

Primer sets for qPCR, ChiP and sequences of siRNA and shRNA can be seen in the table above.
